# Under salt stress quinoa stomatal guard cells control transpiration in an ABA-primed manner

**DOI:** 10.1101/2025.05.24.655927

**Authors:** Shouguang Huang, Maxim Messerer, Heike M. Müller, Sönke Scherzer, M. Rob G. Roelfsema, Christoph Weiste, Markus Krischke, Pamela Korte, Max William Moog, Klaus F. X. Mayer, Peter Ache, Rainer Hedrich

## Abstract

- Stomatal guard cells, located at the interface between the leaf and the atmosphere, play a key role in transpiration control and photosynthetic CO_2_ uptake. Halophytes like *Chenopodium quinoa* tolerate high soil salinity, but the mechanisms governing guard cell responses to salinity stress in relation to the associated epidermal bladder cells (EBCs) remain unknown.
- In this study, responses of *C. quinoa* guard cells under salinity stress and external ABA application were analyzed using RNA profiling and voltage-clamp-based electrophysiology.
- Under salt stress, guard cell RNA profiles reported the activation of ABA synthesis and signaling pathways. However, unlike EBCs, guard cells became transcriptionally insensitive to ABA. Voltage-clamp recordings revealed that under high Na^+^ concentrations, guard cells’ activity of K^+^ uptake channels remained unaffected, while they were impaired in ABA-induced activation of anion channels. As a consequence of a unique guard cell ABA response in salt-adapted plants, stomatal transpiration was reduced and CO_2_ sensitivity enhanced.
- We propose that under salt stress, *C. quinoa* guard cells rewire their hormone signaling to switch from an ABA-sensitive to an ABA-insensitive mode. This adaptation may reflect the halophyte’s ability to perceive salinity as a non-stressful condition, allowing efficient water usage and sustained growth in saline environments.

## Introduction

*Chenopodium quinoa* is a halophytic pseudocereal crop that has unique nutritional value and resistance to multiple abiotic stresses. Because of these valuable traits, quinoa is being cultivated in an ever-increasing number of countries. A major milestone supporting quinoa research was the generation of a high-quality genome draft by Jarvis and colleagues (Jarvis et al. 2017), which provided an essential resource for genetic and molecular studies. In parallel, Zou and colleagues showed that quinoa’s exceptional salt and drought stress tolerance is associated with the expansion of genes involved in ion and nutrient transport, ABA homeostasis, and signaling (Zou et al. 2017).

Quinoa belongs to a group of halophytes that sequester salt and water in giant bladder-shaped epidermal cells (EBCs). EBCs exhibit higher expression of genes related to energy import and ABA biosynthesis compared to the leaf body. Each EBC complex consists of a leaf epidermal cell, a stalk cell, and the bladder. We previously analyzed the molecular mechanisms underlying salt accumulation in bladder cells and demonstrated that, under salt stress, sodium (Na^+^), chloride (Cl^-^), potassium (K^+^), and various metabolites are shuttled from the leaf lamina to the bladders (Bohm et al. 2018). In this process, stalk cells function as both selectivity filters and flux controllers. A unique set of transporters was found to be differentially expressed in the stalk cells and EBCs (Bohm et al. 2018; Bazihizina et al. 2022). Among the genes enriched in the stalk-bladder complex, ion channels, carriers, and sugar transporters were the most pronounced.

Like bladders, stomatal guard cells originate from the specification of epidermal ancestor cells (Lee and Bergmann 2019; Lopez-Anido et al. 2021). Stomata act as water gates, regulating gas exchange between the leaf interior and the atmosphere (Roelfsema and Hedrich 2005). These hydrodynamic valves are powered by changes in the osmotic potential of guard cells (Hedrich 2012). When K^+^ salts accumulate in guard cells, osmotic water influx occurs, leading to an increase in cell volume and turgor pressure. Consequently, the guard cells are pushed apart, causing the stomata to open. During stomatal closure, K^+^ salts are released, leading to water efflux and a reduction in turgor pressure, which closes the stomata. To optimize transpiration, guard cells integrate signals from photosynthesis, the plant’s water status (via ABA), and environmental biotic and abiotic conditions.

Salt can exert its toxic effects on plant biology once it is absorbed from the soil by the roots and transported with the water flow to the shoots and leaves. To analyze the mechanisms of NaCl redistribution in leaves, Grause et al., (2023) directly infiltrated saline solutions into intact tobacco leaves. This acute salt stress caused a reduction in leaf cell pressure, visible as a downward movement of the leaf. However, the leaf fully recovered from the salt challenge, returning to its original position within just 30 to 40 minutes. The injected NaCl diffuses into the cytosol and is eventually transported into the central vacuole. Through this step, the water that is initially lost due to osmosis re-enters the cell, restoring cell turgor and allowing the leaf to stretch again. A similar response has been observed in tobacco plants expressing *Gt*ACR1, a microbe-derived, light-activated anion channel (Ding et al. 2024; Govorunova et al. 2015). Upon light stimulation, leaf cells release anions, leading to membrane depolarization and subsequent K^+^ efflux through GORK-type potassium channels (Huang et al. 2021). As a result of this osmotic loss of solutes, the leaves begin to wilt. However, once the light stimulus is turned off, leaf cells gradually recover, and within approximately 30 minutes, the leaves regain their pre-stimulation turgor level. This dynamic recovery suggests that leaves possess an efficient, ad hoc mechanism for managing acute salt and osmotic stress. Although ordinary leaf cells recover turgor in response to acute 200 mM NaCl stress, the stomatal guard cells seem to reach pre-stress settings only with some delay (Graus et al. 2023). Thus, guard cells seem to handle acute salt stress differently from ordinary leaf cells.

Using the pseudocereal quinoa, we ask: how guard cell function is challenged by salt stress? Incorporating measurements of stomatal transpiration, guard cell electrical properties, and transcription profiles, we provide a comprehensive assessment of the molecular identity and operational modes of major plasma membrane solute transporters, as well as the signaling pathways that regulate them. Our findings demonstrate that under soil salinity, guard cell action is released from ABA signaling control.

## Materials and Methods

### Plant material, growth conditions, and salt treatment

*Chenopodium quinoa* (cv.5020) plants were cultivated in a greenhouse (at 22°C/20°C day/night under a 16h light regime at 300 µmol m^-2^ s^-1^ white light) using common potting soil. NaCl treatment was started after 2 weeks of growth. The NaCl concentration in the irrigation water started at 25 mM NaCl and was increased stepwise by doubling the concentration every second day up to 200 mM or 500 mM. After 5 weeks of salt treatment, plants were used for experiments.

### Infrared gas analysis (IRGA)

Gas exchange measurements were performed on leaves of approximately 7-8 weeks old salt-treated and untreated quinoa plants. Water vapor and CO_2_ concentrations were recorded using a custom-made setup with four parallel water-cooled cuvettes with a gas stream of 0.5 L min^-1^. The leaves were clamped into the cuvettes on the intact plant and an area of 3.8 x 10^-4^ m^2^ each was illuminated at a photon flux density of 300 µmol m^-2^ s^-1^. The gas composition was controlled by mass flow meters (red-y smart series; www.voegtlin.com) as described previously (Muller et al. 2017) and adjusted to 52% RH at 20°C and 0 - 1000 ppm CO_2_. Leaves were illuminated by LEDs (Cree Xlamp CXA2520 LED). The light beams were directed to the cuvettes via fiber-optics (Fiber Illuminator Jumbo; Schott, Mainz, Germany). Recordings were performed with two infrared gas analyzers (LI 7000; Li-Cor, Lincoln, NE, USA).

### ABA-treatment for RNA seq

For ABA stimulation of guard cells, ABA (50 µM ± ABA, Sigma) was fed to the stomata via the petiole of detached leaves, as the dense epidermal bladder growth prevented the usual spray treatment. For comparability, the same procedure was used for ABA application to epidermal bladder cells.

### RNA extraction and sequencing

RNA from quinoa leaves and bladders was extracted, DNase I treated and tested as described (Bohm et al. 2018). Guard cells were enriched using the blender method (Bauer et al. 2013) and RNA was extracted as with bladders. Library preparation and sequencing were performed at the service facility “KFB - Center of Excellence for Fluorescent Bioanalytics” (Regensburg, Germany; www.kfb-regensburg.de).

### ABA quantification

ABA was quantified as described previously (Karimi et al. 2021).

### RNA profiling

The quality of the raw RNAseq reads was analyzed with FastQC (http://www.bioinformatics.babraham.ac.uk/projects/fastqc). The trimming step was performed with Trimmomatic (Bolger et al. 2014) using the parameters ʹILLUMINACLIP:Illumina_PE_adapters.fasta:2:30:10:8:true LEADING:3 TRAILING:3 SLIDINGWINDOW:4:20 MINLEN:60ʹ. The reads were mapped to the quinoa reference transcriptome (Jarvis et al. 2017) using Kallisto (Bray et al. 2016). Differentially expressed genes (DEGs) were calculated using EdgeR with a significance threshold of p<0.05 (Robinson et al. 2010). RNAseq data of GCs ± salt and ± ABA were submitted to EMBL-EBI (https://www.ebi.ac.uk/biostudies) under the accession number E-MTAB-14806. The EBC ± ABA data can be found under the accession E-MTAB-10419.

Table S1 (GCs ± salt and ± ABA) was generated by removing all non-expressed DEGs that had (transcripts per million) TPM<1 in all treatments simultaneously. Likewise, we also removed all genes that were not differentially expressed in any of the contrasts (i.e. simultaneously empty cells in all comparisons, or value = 0).

### Electrophysiological recordings on guard cells

Electrophysiological recordings were performed on guard cells in intact leaves, as described previously (Huang et al. 2024). The adaxial side of the leaves was gently fixed onto a Petri dish using double-sided adhesive tape and incubated in a bath solution (50 mM KCl, 1 mM CaCl₂, and 5 mM MES/BTP, pH 6, unless otherwise specified). To improve the visualization of guard cells, the leaf surface was gently brushed several times with a soft paintbrush to reduce its hydrophilic properties. The Petri dish containing the samples was then placed on a microscopic table (Axioskop 2FS, Zeiss, Germany) for four hours before measurement. Guard cells were impaled with double-barreled microelectrodes made from borosilicate glass capillaries (inner/outer diameter = 0.56/1.0 mm; Hilgenberg, Germany). These capillaries were aligned, heated, twisted 360 degrees, and pre-pulled using a vertical puller (L/M-3P-A, Heka, Germany), followed by a final pull on a horizontal laser puller (P2000, Sutter Instruments, CA, USA). The electrodes were filled with 300 mM KCl and had a tip resistance ranging from 130 to 180 MΩ. The electrodes were mounted into the holder of a piezo-driven micromanipulator (MM3A, Kleindiek, Reutlingen, Germany), which was used to impale single guard cells. Both barrels of the microelectrode were connected via Ag/AgCl half-cells to headstages with an input impedance of 100 GΩ. The headstages were further linked to a custom-made amplifier (Ulliclamp01) equipped with an internal differential amplifier for voltage clamp measurements. Electrical signals were low-pass filtered at 0.5 kHz using a dual low-pass Bessel filter (LPF 202A; Warner Instruments Corp., USA) and recorded at 1 kHz with an interface (USB-6002, NI, USA) controlled by WinWCP software (Dempster 1997). To load sodium, malate, or acetate into guard cells, the tips of the electrodes were filled with NaCl, potassium malate, or potassium acetate, as specified in the figure legends.

### Stomatal aperture assay

Stomatal aperture assays were performed on stomata located on the abaxial side of leaves, prepared similarly to the impalement experiment. Stomata were visualized using an immersion objective (W Plan-Apochromat, 63×; Zeiss) mounted on an upright microscope (Axioskop 2FS; Zeiss, Jena, Germany). The microscope was equipped with a sCMOS camera (Prime BSI Scientific CMOS, Teledyne Photometrics), which was controlled by VISIVIEW software (Vistron, Germany). The same stoma was monitored before and after ABA application over a 30-minute time course. The concentration of the applied ABA is specified in the figure legend. Stomatal apertures were quantified using FIJI/IMAGEJ software.

### Ion flux measurements

Anion flux was measured in proximity to stomatal pores in epidermal strips using non-invasive Scanning Ion-Selective Electrodes (SISE), as previously described (Huang et al. 2024; Ahmad et al. 2025). For this purpose, epidermal strips were gently peeled and fixed onto a Petri dish (diameter: 3 cm), which was then filled with 1.5 ml of bath solution (0.1 mM KCl, 0.1 mM KNO₃, 0.1 mM CaCl₂, 0.1 mM MES/BTP, pH 6). The samples were kept in the dark for at least four hours prior to measurement. Electrodes were pulled from borosilicate glass capillaries without filaments (diameter: 1.0 mm, Science Products, Hofheim, Germany) using a vertical puller (Narishige, Tokyo, Japan), baked overnight at 220 °C, and subsequently silanized with N,N-Dimethyltrimethylsilylamine (Sigma-Aldrich). The silanized electrodes were backfilled with 500 mM NaCl and chloride ionophore I cocktail A (Sigma-Aldrich). Only electrodes that exhibited a shift of more than 58 mV per 10-fold ion concentration change were used for measurements. The electrode was connected to the headstage of a custom-built microelectrode amplifier via Ag/AgCl half-cells and positioned approximately 5 μm from a stoma using a micromanipulator (PatchStar, Scientifica, Uckfield, UK). The electrodes were moved over a distance of 50 μm at an angle of 45° to the sample surface, with measurements taken at 10-second intervals. Raw data were acquired using an interface (USB 6002, NI, USA) controlled by custom-built LabVIEW-based software, Ion Flux Monitor. Offline, raw voltage data were converted into ion flux data using the custom-made Ion Flux Analyzer program, as described (Ahmad et al. 2025).

### Statistics

Statistical analysis was performed using Origin 2021b software. Data normality was determined by Shapiro-Wilk test. If not stated otherwise, normally distributed data was analyzed using parametric tests (two-tailed Student’s *t*-test for groupwise comparisons and one-way ANOVA with Tukey post-hoc test for multiple comparisons). In case individual data sets were not normally distributed non-parametric tests were applied (Mann-Whitney *U* test for groupwise comparisons and Kruskal-Wallis ANOVA with Dunns post-hoc test for multiple comparisons). For reasons of clarity statistics for large scale datasets are only provided in Supplemental **Table S3**.

## Results

### Salt exposure enhances water use efficiency

Stomata are microscopic pores that control transpiration. By adjusting the stomatal aperture, plants can optimize CO_2_ uptake and water loss of the leaves. Transpiration thus serves as proper read-out for the degree of stomatal opening. To study the response of stomatal guard cells to soil salinity, quinoa plants were exposed to 200 or 500 mM NaCl for five weeks. Infrared gas analysis (IRGA) was applied to monitor transpiration and CO_2_-assimilation in leaves on intact plants. In well-watered, NaCl-free conditions, dark transpiration of cultured plants can be quite substantial (Lu and Fricke 2023). Quinoa plants grown on salt-free soils were not found to be an exception. Background transpiration of plants exposed to 200 and 500 mM NaCl was reduced by 60 % compared to the controls (**Fig. 1a**). This indicates that, in darkness and hence absence of photosynthetic CO_2_ fixation, salt-stressed quinoa plants save water by reducing transpiration. In the light transpiration increased under all conditions, reflecting that the stomata remained light-sensitive despite high salt exposure. In control plants, a steady state was reached approximately 100 min after illumination onset. Under 200 mM NaCl, stomatal opening reached a plateau at 33% of the control level after 70 min, while at 500 mM NaCl, it reached steady state at 25% of the control, in less than 30 min.

**Figure 1.**
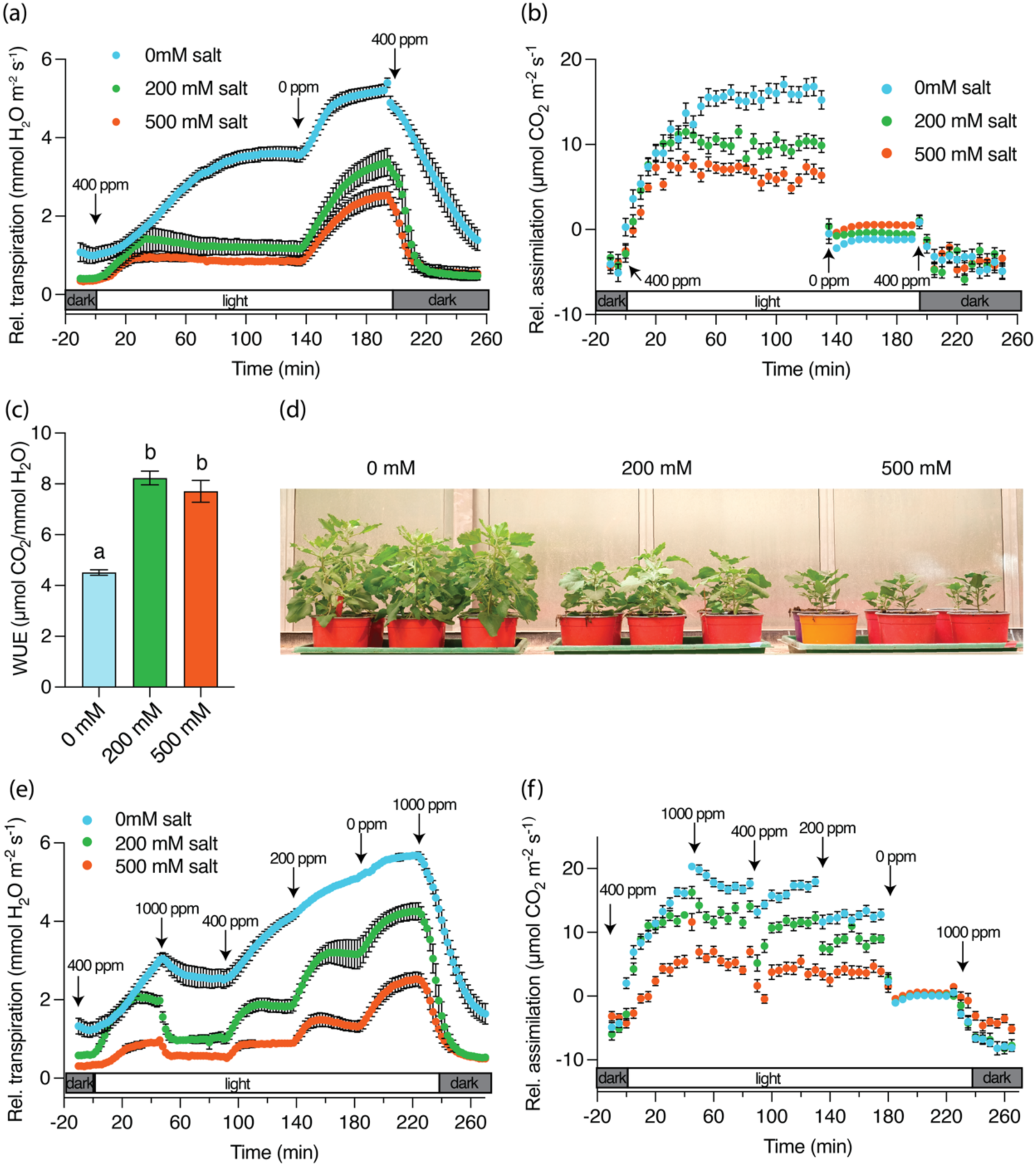
Long-term salt exposure increases water use efficiency (WUE) in Quinoa. Plants were treated long-term with 0 mM (control), 200 mM, or 500 mM NaCl. Gas exchange measurements began in darkness at an ambient CO₂ concentration (400 ppm). At time t = 0, light was switched on (300 µmol m^-2^ s^-1^ PAR). After 140 minutes, once steady-state transpiration and CO_2_ uptake were reached, CO_2_-free air was supplied. After an additional 60 minutes, 400 ppm CO_2_ was reintroduced, and light was turned off. **(a)** Relative transpiration rate, averaged over 2-minute intervals (n = 8, mean ± SE). **(b)** Relative CO_2_ assimilation rate, averaged over 5-minute intervals (n = 8, mean ± SE). **(c)** Intrinsic WUE, calculated as the ratio of assimilation to transpiration, averaged over 30 minutes (minutes 105–134) under light and 400 ppm CO_2_ (n = 8, mean ± SE). **(d)** Phenotype of the quinoa under different salt treatments. NaCl treatment was initiated after 2 weeks of growth. The NaCl concentration in the irrigation water started at 25 mM and was increased stepwise by doubling every second day until reaching the final concentrations of 200 mM or 500 mM. Photographs of plant phenotypes were taken 5 weeks after the start of salt treatment. **(e -f)** Plants were long-term treated with 0 mM (control), 200 mM, or 500 mM NaCl. Gas exchange measurements began in darkness at ambient CO_2_ (400 ppm). After 45 minutes, light was switched on (time t = 0). Every subsequent 45 minutes, CO_2_ concentration was changed sequentially to 1000 ppm, 400 ppm, 200 ppm, and 0 ppm. In the final step, CO_2_ was increased again to 1000 ppm while light was simultaneously switched off. (e) Relative transpiration rate, averaged over 2-minute intervals (n = 8, mean ± SE). (f) Relative CO_2_ assimilation rate, averaged over 5-minute intervals (n = 33– 40, mean ± SE). Statistically significant differences were determined by one-way ANOVA followed by Tukey’s post hoc test (see Table S3).

CO_2_-depletion of the air triggered stomatal opening at all conditions **(Fig. 1a)**, which revealed that stomata remain CO2 sensitive after salt treatment. Nevertheless, the degree of stomatal opening at 400 ppm was strongly reduced after salt treatment. When normalized to the transpiration maximum under CO_2_-free air, control plants transpired up to 70% at 400 ppm CO_2_, whereas plants exposed to 200 and 500 mM NaCl showed only 35% of this maximum **(Fig. 1a)**. Simultaneously, CO_2_ assimilation decreased to 60% of the control level at 200 mM NaCl and to 44% at 500 mM **(Fig. 1b)**, showing that CO_2_ assimilation is less affected than transpiration. As a result, water-use efficiency (WUE; **Fig. 1c**) increased by 80% under 200 mM salt and by 60% under 500 mM salt. Despite the improved WUE, reduced CO_2_ assimilation was reflected in growth performance. Plants treated with 200 mM NaCl reached 54% of the height of untreated controls, and those under 500 mM reached only 33% **(Fig. 1d)**. However, given that plants still appeared healthy even under 500 mM NaCl **(Fig. 1d)**, the enhanced WUE likely represents a key factor contributing to quinoa’s salt tolerance.

To answer the question of whether salt treatment affects the CO_2_ sensitivity of stomatal action, we challenged quinoa stomata with different CO_2_ concentrations. In IRGA measurements, stomata were first allowed to open in the light at ambient CO_2_ (400 ppm), then exposed sequentially to 1000 ppm, 400 ppm, 200 ppm, and finally CO_2_-free air (**Fig. 1e**). Control plants exhibited significantly higher transpiration at all CO_2_ concentrations compared to plants treated with 200 mM NaCl, which in turn transpired more than those treated with 500 mM NaCl. At 1000 ppm CO_2_, transpiration of control plants in the light dropped only marginally, whereas in plants treated with 200 or 500 mM NaCl, transpiration nearly returned to the dark baseline level. This indicates that quinoa stomata under salt exposure profit from increased CO_2_ sensitivity. This is exemplified by the fact that the CO_2_ assimilation at 1000 and 400 ppm CO_2_ under all salt conditions remained unchanged (**Fig. 1f**).

The CO_2_ sensitivity of stomata has been shown to be ABA-dependent, with the ABA receptors PYR1 and PYL2 playing key roles in Arabidopsis guard cells (Dittrich et al. 2019; Chater et al. 2015). We therefore asked whether salt stress feeds back on ABA levels and thereby influences ABA-triggered stomatal action. Surprisingly, we found that under control conditions, the ABA content in quinoa leaves was nearly 10 times higher than in the glycophyte *Arabidopsis thaliana* and the halophyte *Thellungiella salsuginea* (Karimi et al. 2021). Leaf ABA content increased to 1.7-fold of the control at 100 mM NaCl and reached approximately threefold at 200 mM, but did not rise further with higher salt concentrations (300, 400, or 500 mM; **Fig. S1a**). In contrast, guard cell ABA levels remained unchanged up to 200 mM NaCl, then began to rise and peaked at 400 mM (**Fig. S1b**). This trend aligns with the salt-dependent reduction in stomatal transpiration.

### Salt stress activates a substantial population of ABA-responsive genes

To gain insight into the impact of salt treatment on the molecular mechanism of guard cell ABA responses, we used a transcriptomic approach (Bauer et al. 2013). Epidermal samples containing only viable guard cells were collected from i) salt-stressed and control plants and those ii) additionally treated with ABA, and were subjected to RNA sequencing. In the RNA-seq data, when comparing the top 20 most abundant genes identified in the earlier study with our dataset from salt-treated and control guard cells (Rasouli et al. 2022), we observed a 75% overlap in differential expression. Interestingly, the transcriptome of our salt- and ABA-treated guard cells showed almost 100% correlation with the dataset of salt-only treated guard cells reported by (Rasouli et al. 2022). To explore this further, we performed a more detailed analysis of the ABA-responsive gene set.

Salt treatment alone resulted in almost 10,000 differentially expressed genes (DEGs, **Fig. 2a, Table S1a**), ABA treatment without salt resulted in 4,000 DEGs (**Fig. 2a, Table S1b**), of which almost 2,600 were also affected by salt without ABA (**Fig. 2a, Table S1d**). This indicated that salt alone triggers ABA synthesis and signal transduction. Under salt stress, only 2,200 genes were regulated by additional ABA treatment (**Fig. 2a, Table S1c**). Of these 2,200, 1,400 (64%) were also affected by salt treatment alone. Interestingly, 90% of these DEGs were completely antagonistically regulated (**Fig. 2b, Table S1e**). This population thus represents genes for which the ABA effect is remarkably stronger than that of salt. Among them, we found genes related to the hormones ABA (5 genes), ethylene (8 genes), and jasmonic acid (6 genes), as well as 5 cell wall-modifying expansins and 7 transporters (see below).

**Figure 2.**
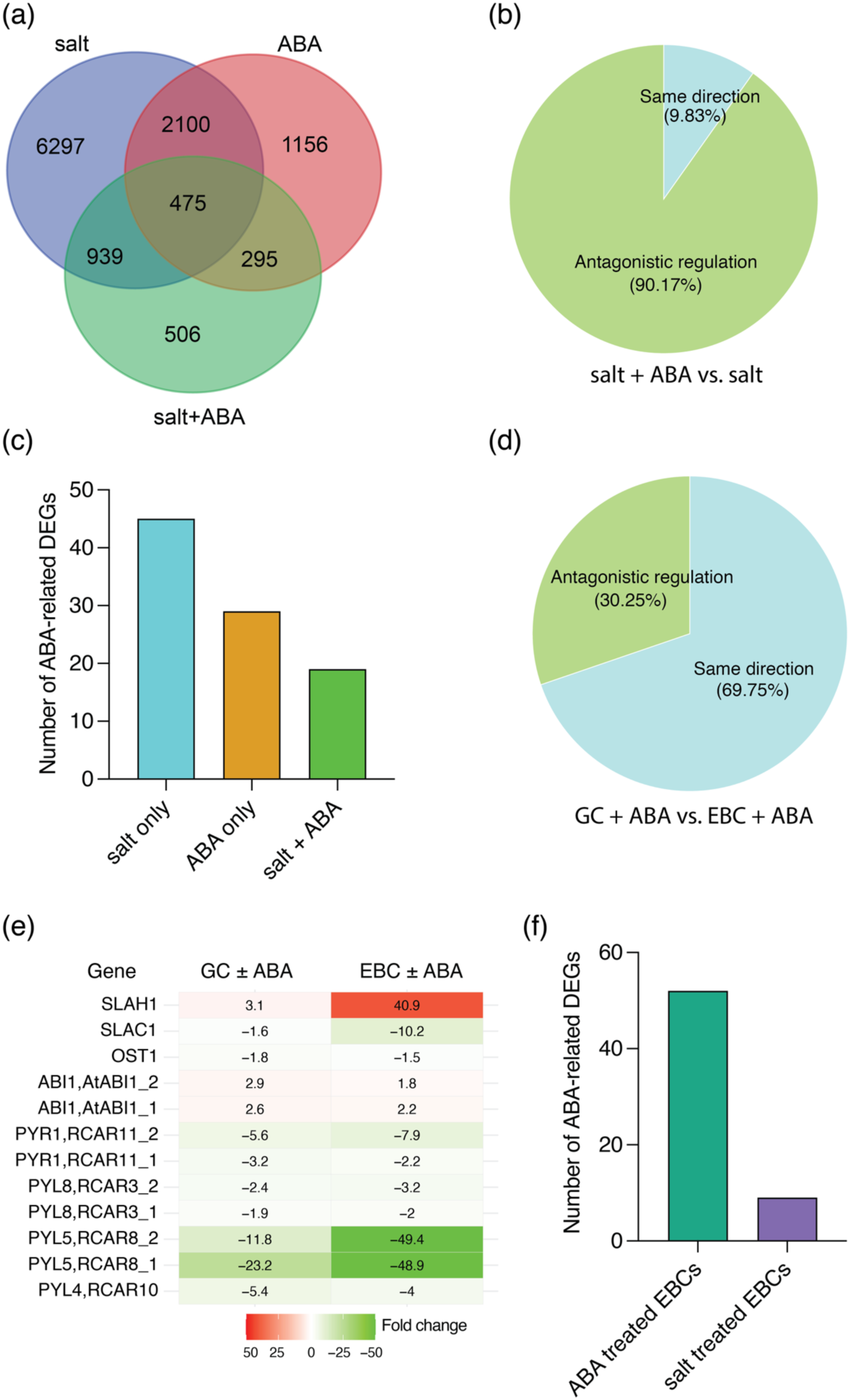
Transcriptomic comparison of salt and ABA responses in quinoa guard cells and epidermal bladder cells (EBCs). **(a)** Venn diagram showing the overlap of differentially expressed genes (DEGs) in guard cells exposed to salt, ABA, or combined salt + ABA treatments. **(b)** Pie chart illustrating that 90% of the 1,400 DEGs shared between salt + ABA and salt-only treatments in guard cells are regulated in opposite directions. **(c)** Bar plot comparing the number of ABA-signaling-pathway-related DEGs under salt-only, ABA-only, and salt + ABA conditions, with salt treatment inducing the highest number. **(d)** Pie chart indicating that 70% of the 1,382 DEGs shared between guard cells and EBCs in response to ABA are regulated in the same direction. **(e)** Heatmap showing fold changes in selected ABA signaling genes following ABA treatment in guard cells and EBCs. **(f)** Bar plot showing the number of ABA-related metabolic pathway DEGs in EBCs following ABA (52 genes) and salt treatment (9 genes).

In the datasets of ABA treatments with and without salt, we found the lowest overlap—only 770 genes (**Fig. 2a, Table S1f**)—suggesting that these genes are specifically regulated by ABA, independent of salt stress immanent factors. Among genes involved in ABA synthesis, conjugation/degradation, and signaling, most DEGs were again found in the salt-treated fraction (45 DEGs, **Table S1a1**), supporting the conclusion that the ABA signaling pathway is already activated by salt stress. In contrast, significantly fewer DEGs were detected in the ± ABA treatment without salt (**Fig. 2c**, 29 DEGs, **Table S1b1**). It is well known that high ABA levels can reduce further ABA synthesis, which is reflected in our data by the strong downregulation of ABA receptors and concurrent induction of PP2Cs and ABA-conjugating enzymes (Dittrich et al. 2019). The lowest number of ABA-related DEGs was found in the comparison between ABA-treated and control samples under salt stress (**Fig. 2c**, 19 DEGs, **Table S1c1**). Under these conditions, almost all ABA-driven gene regulations appear to have already been triggered by the salt-induced ABA increase.

Guard cells and epidermal bladder cells (EBCs) both reside in the epidermis. We therefore asked whether both cell types share the same ABA sensitivity under salt stress. To generate a bladder-specific dataset, ABA was fed via the petiole to excised leaves, and RNA was sequenced from isolated EBCs (**Table S2**). When comparing the ABA responses of guard cells and EBCs, their transcriptional responses to ± ABA were quite similar (**Table S2a**). We found 1382 co-regulated DEGs in the comparison of EBCs and guard cells ± ABA. Of these, 70% were regulated in the same direction and 30% in opposite directions (**Fig. 2d**). This indicates that these two cell types respond similarly to ABA. This fact is exemplified by ABA receptors (PYL/RCAR), which are downregulated under ABA in both EBCs and guard cells (**Fig. 2e**) (Dittrich et al. 2019). At the same time, several PP2Cs (e.g. ABI1), OST1, and anion release channels of the SLAC/SLAH type are regulated in the same direction (**Fig. 2e**); ABA perception and signaling thus is supported in both systems, underlining the notion that the bladders under ABA behave like guard cells (**Table S2b**). The correlation was, however, found to be less significant when comparing EBCs ± ABA and guard cells ± salt, although the total number of commonly regulated genes was higher (3238 in total, but only 52 % in the same direction and 48 % in opposite directions). At a first glance it looked like several transporters in ABA-treated EBCs and only salt-stressed guard cells went in the same direction (**Table S2c**). However, analysis of the ABA-related metabolic pathways revealed major differences between the two cell types. We compared the EBC ± ABA dataset with previously published EBC transcriptomes before and after salt treatment (Bohm et al. 2018) (**Table S2d**). ABA treatment induced 52 DEGs in EBCs (**Fig. 2f, Table S2b**), but salt treatment alone triggered only 9 DEGs **(Fig. 2f)**. These results demonstrate that EBCs, unlike guard cells, cannot initiate the ABA signaling pathway in response to salt stress alone.

### Guard cell K^+^ channels’ performance is maintained under salt stress

Salt exposure increases quinoa WUE most likely via ABA-dependent stomata regulation. Stomatal movement is mediated via guard cell ion transporters. Na^+^ and K^+^ have similar physical properties, and one may thus expect an impact of Na^+^ on K^+^ channels. However, so far guard cell plasma membrane endogenous electrical properties of the intact plants grown under salt stress have not yet been tested in any species, including the model *Arabidopsis thaliana*.

To bridge this gap, we tested the properties and capacity of the K^+^- and anion-driven osmotic motor of the guard cells under salt stress. Guard cell ion transport was analyzed in two different scenarios: i) direct NaCl loading into guard cells or ii) indirect NaCl exposure via increasing soil salinity affecting the roots. In both situations, stomatal guard cells in intact leaves were impaled with double-barreled electrodes to monitor their electrical fingerprint. Under voltage-clamp conditions, the membrane potential was altered in 20 mV steps from - 100 mV to potentials ranging from +20 mV to -200 mV. When the guard cell extracellular medium (bath) contained 1 mM K^+^ only, pronounced outward currents of up to about 2000 pA at 0 mV were recorded (**Fig. 3a**). Under such low K^+^ conditions, inward currents were of small amplitude only. Upon the rise in external K^+^ concentration to 10 mM in the first step and 50 mM in the second, outward currents dropped to 1000 pA, while inward currents at -200 mV increased to 1000 pA (**Fig. 3a-b**). When superimposed, the I/V and open-probability curves shifted in K^+^- and voltage-dependent manner. This behavior together with the change in reversal potential by 70 mV identifies both the outward and inward rectifying currents mediated by K^+^ selective KAT1/2- and GORK-type K^+^ channels (Hedrich 2012).

**Figure 3.**
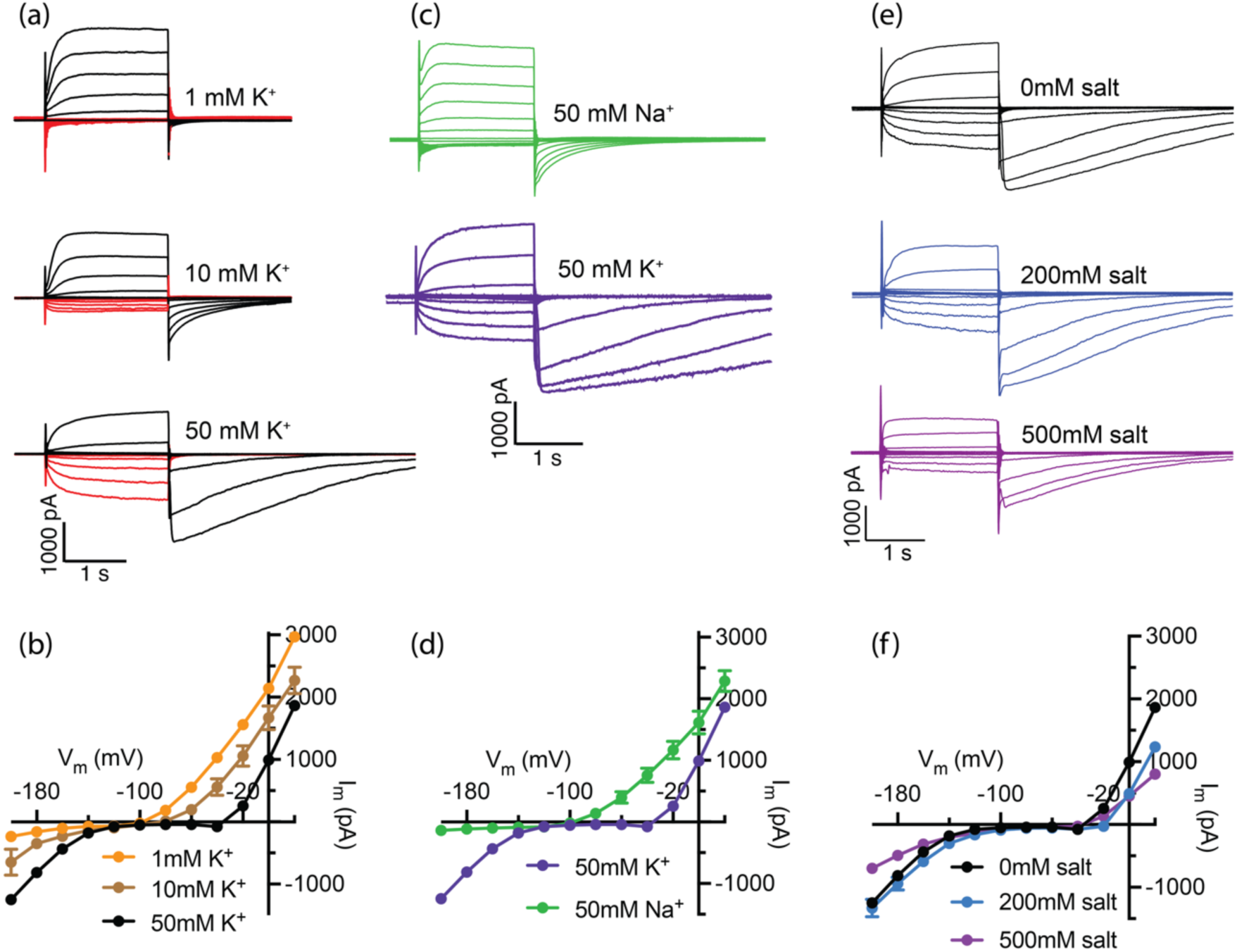
Effects of salt stress on quinoa guard cell K^+^ channel activity. **(a)** Representative current traces showing outward (black) and inward (red) K^+^ currents recorded in bath solutions containing 1, 10, or 50 mM KCl. Guard cells in intact leaves were impaled with double-barreled electrodes filled with 300 mM KCl. Cells were voltage-clamped at a holding potential of –100 mV and stimulated with 2 s pulses to test potentials ranging from +20 mV to –200 mV in 20 mV increments. **(b)** Current–voltage relationships of steady-state currents recorded in 1, 10, and 50 mM KCl (n = 6, 7, and 8, respectively). **(c)** Representative current traces recorded in bath solutions containing either 50 mM NaCl or 50 mM KCl. Inward rectifying currents were strongly activated by hyperpolarization (–120 to – 200 mV) in KCl but not in NaCl, indicating Na^+^ does not substitute for K^+^ in activating inward K^+^ channels. **(d)** Steady-state current–voltage relationships under the same conditions as in (c). **(e)** Representative K^+^ current traces recorded in guard cells from plants treated with 0, 200, or 500 mM NaCl. **(f)** Current–voltage relationships of steady-state K^+^ currents under each treatment condition as in (e).

To challenge guard cells with a sudden salt load, we replaced 50 mM KCl with an equal amount of NaCl. Upon replacement of K^+^ by Na^+^, activation of outward K^+^ currents was already observed at -100 mV, whereas these channel open only at -20 mV for the KCl controls (**Fig. 3c-d**). The reversal potential was shifted from -40 to -140 mV (**Fig. 3d**). This shift in voltage-dependence to more hyperpolarized membrane potentials was associated with an increase in activation kinetics of GORK-type channels (**Fig. 3c**). Thus, in the presence of Na^+^, the guard cell outward rectifier is responding like a K^+^ selective channel facing low extracellular K^+^. The response of the inward rectifier to the K^+^/Na^+^ exchange was even more pronounced (**Fig. 3c - d**). In line with the nominal loss of the K^+^ substrate, the inward rectifier became electrically silent.

Next, we asked the question of how the guard cell plasma membrane ion channels characterized above are affected when quinoa plants are cultured in up-salted soils (see methods). When glycophyte crops are faced with soil salinity of 200 mM NaCl, they do not survive (Munns and Gilliham 2015). Under such conditions, quinoa growth and development were not too different from salt-free controls (**Fig. 1d**). Likewise, guard cell currents monitored in the presence of 50 mM extracellular KCl were similar (**Fig. 3e-f**). This situation did not change qualitatively when plants were cultured on 500 mM NaCl. However, inward and outward currents at -200 and 0 mV were reduced by 44% and 54%, respectively. Additionally, the activation kinetics of K^+^ channels were accelerated (**Fig. 3e**). This may mean that even in 500 mM NaCl the cytosolic Na^+^ does not reach a level high enough to inhibit the K^+^ outward rectifier (Thiel and Blatt 1991). To test how the quinoa K^+^ outward channels respond to elevated cytosolic Na^+^ concentrations, we tip-loaded microelectrodes with 50 mM NaCl before guard cell insertion. Under these conditions, K^+^ outward currents were found largely suppressed (**Fig. S2**). Together, these findings indicate that in salt-treated plants, Na^+^ concentration in the guard cell cytoplasm is not reaching a level that can terminate K^+^ release. Thus, guard cells of plants under high salt appear to have ways to prevent cytosolic Na^+^ from reaching levels high enough to block GORK-like depolarization-activated K^+^ channels.

In addition to K^+^-dependent changes in inward and outward current amplitudes, we observed slowly deactivating tail currents following depolarizing pulses. Both the amplitude and duration of these tail currents increased with external K^+^ concentration (**Fig. 3**), resembling depolarization-activated S- and R-type anion currents(Schroeder and Keller 1992; Keller et al. 1989). Replacing intracellular Cl^-^ with malate or acetate did not affect the voltage dependence of K^+^ channels **(Fig. S3a - b)**, but reduced tail current duration—more so with acetate than malate **(Fig. S3c - d)** (Imes et al. 2013; Meyer et al. 2010; Pei et al. 1997; Blatt 1987). These results suggest that the tail currents reflect combined activity of GORK-type outward K^+^ channels and R-/S-type anion channels.

### Severe salt stress reduces ABA sensitivity of both anion channels and stomatal closure

In the following, we asked how salt stress addresses anion channel activity in guard cells. To this end, anion fluxes were measured using the Scanning Ion-Selective Electrode (SISE) technique (Ahmad et al. 2025). Epidermal strips from plants grown under 0, 200, and 500 mM NaCl were used to compare anion fluxes in guard cells. After equilibration in a standard bath solution, no measurable anion fluxes were detected in control plants or those grown under 200 mM NaCl (**Fig. 4a - b**). In contrast, plants subjected to high salt stress (500 mM NaCl) exhibited a small anion efflux (**Fig. 4c and d**). To assess whether this effect is related to ABA signaling, guard cells were exposed to 100 µM ABA. ABA treatment triggered a pronounced anion efflux in control plants (**Fig. 5a**). However, the ABA-induced efflux measured 5 minutes after hormone application was slightly reduced in plants grown under 200 mM NaCl compared to the control (**Fig. 4a - b**). Notably, in plants grown under 500 mM NaCl, ABA failed to further enhance the already elevated anion efflux (**Fig. 4c - d**), suggesting desensitization of the ABA response.

**Figure 4.**
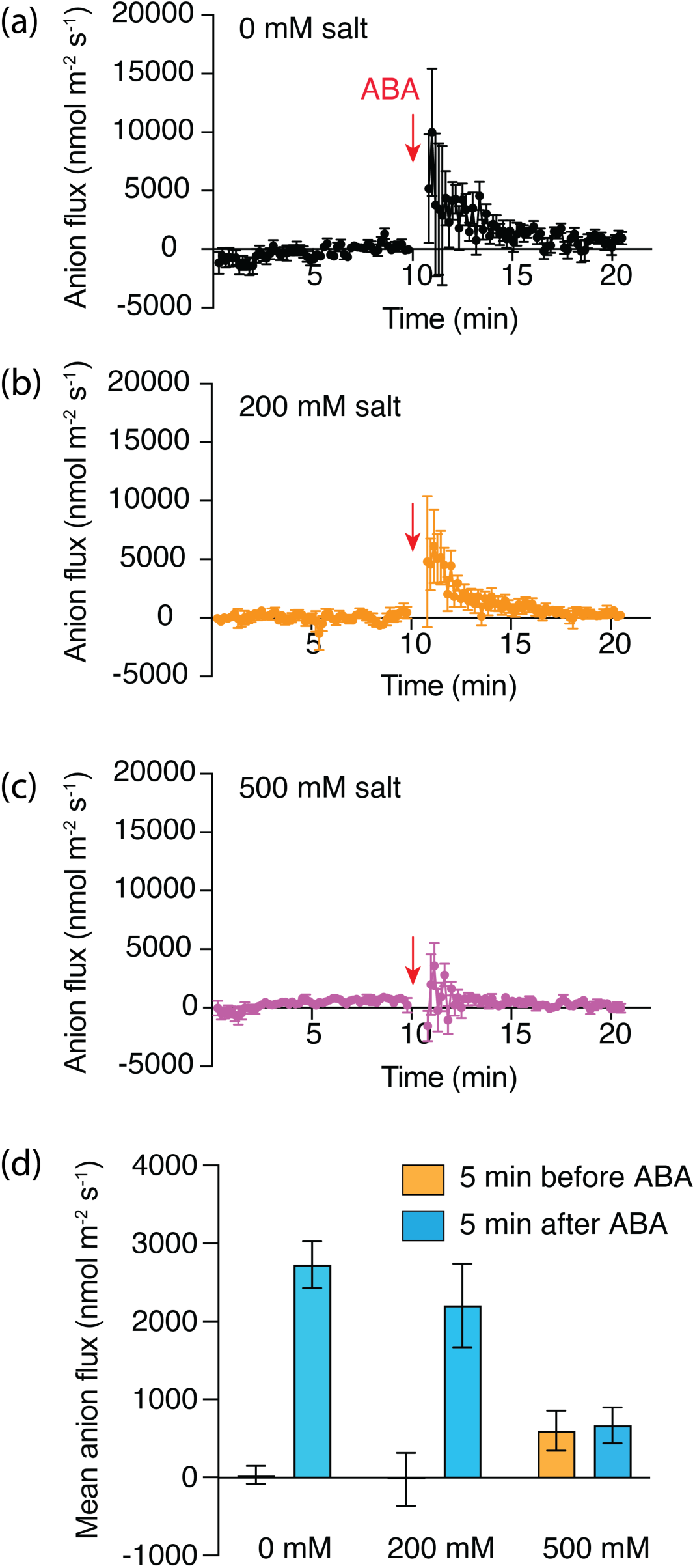
ABA induces anion efflux in quinoa guard cells under different salt-growth conditions. **(a-c)** Anion efflux from quinoa guard cells was measured using the Scanning Ion-Selective Electrode (SISE) technique. Plants were grown under control, 200 mM NaCl, or 500 mM NaCl conditions as indicated. Application of 100 µM ABA at 11 minutes triggered an immediate and transient anion efflux in guard cells from control (0 mM salt, black) and 200 mM NaCl-grown plants (brown), but not in plants grown under high salt stress (500 mM NaCl; purple). Data are shown as mean ± SE (n = 15, 18, and 24, respectively). **(d)** Mean anion efflux measured 5 minutes before and after ABA application for each treatment condition.

**Figure 5.**
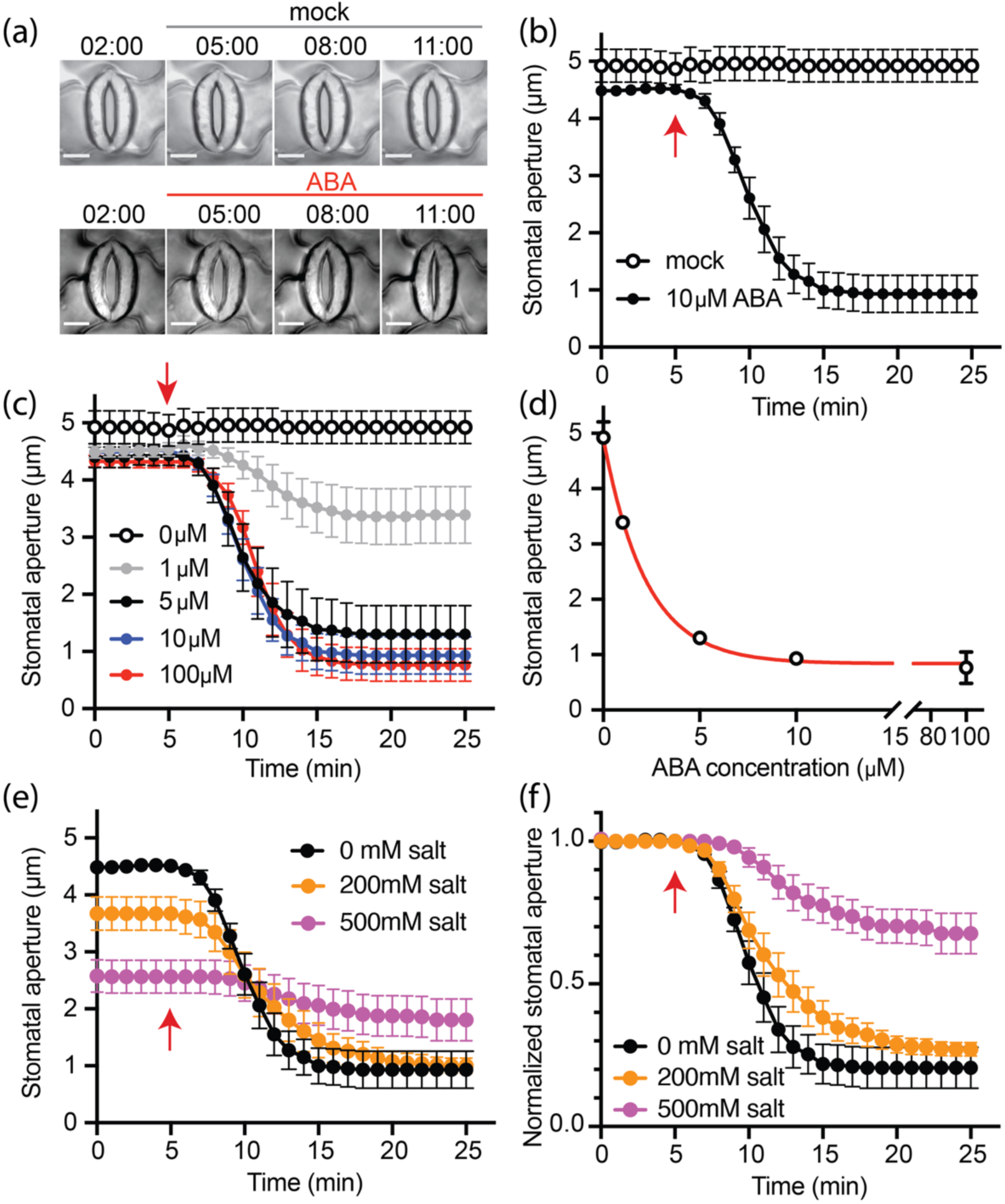
Stomatal responses of quinoa to ABA. **(a)** Representative images of stomata in quinoa leaves before and after mock treatment (upper panel) or 10 µM ABA treatment (lower panel). Time points correspond to the time axis in (b). Scale bar: 10 µm. See also videos S1-S2. **(b)** Time course of stomatal aperture changes following ABA application. The arrow indicates the time of ABA addition to the bath solution. **(c - d)** Dose dependency of ABA-induced stomatal closure. Data were fitted with an exponential function (red curve in d). **(e)** Stomatal response to 10 µM ABA in control plants and those grown under 200 mM or 500 mM NaCl. **(f)** Normalized stomatal aperture from (e) following ABA treatment.

Together, these data indicate that guard cell K^+^ and anion currents are only marginally affected by moderate soil salinity. However, under high salinity (500 mM NaCl), ABA sensitivity of anion efflux is lost. Given that anion channels are key for stomatal closure (Huang et al. 2021), we next monitored stomatal aperture changes in response to ABA in plants grown at 0, 200, and 500 mM NaCl. Stomatal opening was induced by microscope light, and ABA was applied directly to the bath solution. Upon onset of a 10 µM stimulus, stomata of the controls closed in about 10 min (**Fig. 5a-b** and **videos S1-S2**). This response was both time- and ABA concentration-dependent. When guard cells were challenged by 1, 5, 10, and 100 µM extracellular ABA, the resulting dose–response curve was well fitted by an exponential function. The half-maximal stomatal closure was reached at an ABA apparent concentration of 1.54 µM (**Fig. 5c-d**). Compared to control plants, stomatal apertures were less open in plants treated with 200 mM salt, and they further reduced in size under higher salt concentrations (**Fig. 5e**). In line with anion flux measurements, the stomatal response to ABA in 200 mM salt-treated plants was comparable to that of control plants but was significantly impaired in plants exposed to higher salt levels (**Fig. 5e**). Specifically, ABA-induced stomatal aperture reduction was only 30% in 500 mM NaCl-treated plants, compared to 80% in control plants (**Fig. 5f**). Moreover, the rate of stomatal closure decreased from 0.12 mm/min in control plants to 0.036 mm/min in plants exposed to 500 mM NaCl (**Fig. 5f**).

## Discussion

### Role of ABA in salt stress responses of quinoa

High loads of salt in the soil hamper the growth of most crop plants, but quinoa has found ways to cope with this challenge. This means that quinoa can overcome the toxicity of Na^+^ and Cl^-^, as well as the high soil osmolarity that restricts water uptake. Our data reveal that the high soil osmolarity has a major impact on stomatal movements in quinoa, via an increase of the ABA level in leaves **(Fig. S1a)**, as well as guard cells **(Fig. S1b)**. These higher ABA levels are likely to restrict stomatal opening **(Fig. 5e)** and thus results in a lower stomatal conductance of quinoa plants that are exposed to 200 and 500 mM NaCl **(Fig. 1a)**. However, the increases in ABA level are moderate and mainly reduce the dark transpiration, but do not inhibit stomatal opening in light, or at low atmospheric CO_2_ concentration **(Fig. 1)**.

Just as in other plant species (Huang et al. 2019; Roelfsema et al. 2004; Guo et al. 2023), ABA activates anion channels in quinoa guard cells **(Fig. 4)**, which provokes anion efflux and leads to stomatal closure. However, salt stress seems to cause only a moderate increase in the ABA level of guard cells, since stimulation of the cells with ABA can still induce anion efflux **(Fig. 4b)** and provoke closure of stomata **(Fig. 5e-f)**. Quinoa just seems to fine-tune its ABA responses to reduce the stomatal conductance and enhance the water use efficiency, but the stomata are not fully closed and remain sensitive to light and CO_2_.

### Impact of salt stress and ABA on gene transcription

Salt stress not only limits the opening of quinoa stomata, but also provokes a wide range of changes in the transcriptome of leaves, epidermal bladder cells and guard cells. In line with the increase in ABA concentration during salt treatment (**Fig. S1**), a pronounced overlap was found between DEGs induced by salt and ABA fed through the petiole of leaves (**Fig. 2a**). To our surprise, the transcript level of a subset of 700 genes was affected by salt treatment, but still sensitive to ABA application. Possibly, the salt treatment led to a more gradual and thus less strong ABA response, as the sudden increase of the ABA level imposed by petiole feeding. Alternatively, salt treatment may repress ABA-regulation of a subset of genes, via an ABA-independent mechanism. Further experiments focusing on the salt-induced ABA-dependent and independent pathways will be required to answer the open question.

Despite their functional differences, guard cells and EBCs display notably similar transcriptional responses to exogenous ABA. Key genes involved in ABA-induced stomatal closure—such as PYL/RCAR receptors, clade A PP2Cs, OST1, and SLAC1/SLAH3 anion channels—are consistently regulated in both cell types following ABA treatment (**Fig. 2e**), indicating that EBCs, like guard cells, possess a functional ABA signaling machinery and respond to ABA at the transcriptomic level.

In contrast, salt stress triggers markedly different transcriptional responses in guard cells and EBCs, suggesting distinct signaling mechanisms. In guard cells, ABA- and salt-responsive genes are regulated in a largely consistent manner, indicating that salt stress stimulates local ABA biosynthesis (**Fig. S1**) and activates downstream signaling similar to drought-induced responses. However, in EBCs, the expression of the same genes is often regulated in opposite directions under salt versus ABA treatment, implying that salt does not activate the ABA pathway in EBCs. This is supported by transcriptomic comparisons: ABA treatment induces 52 ABA-related DEGs in EBCs, whereas salt alone affects only 9 genes (**Fig. 2f**). These findings suggest that EBCs, unlike guard cells, cannot initiate the ABA signaling pathway in response to salt stress alone. While salt induces guard cell–intrinsic ABA synthesis (**Fig. S1**), EBCs seem to rely on exogenous ABA supplied via systemic drought signaling pathways. This difference may reflect the role of EBCs as osmotically buffered compartments—possibly functioning as water reservoirs during drought, but not perceiving progressive salinity as a stressor. However, there are contradictory evidence for this reservoir hypothesis (Moog et al. 2023; Adolf et al. 2013; Shabala and Mackay 2011; Agarie et al. 2007), and further research into this EBC function is needed.

### Guard cell ion channels and salt stress

The physical properties of Na^+^ are similar to K^+^, and early experiments with guard cells were therefore focused on the direct impact of Na^+^ on the properties of K^+^ channels (Thiel and Blatt 1991; Véry et al. 1998). These studies showed that cytosolic Na^+^ inhibits GORK-like channels, just as found for guard cells of quinoa **(Fig. S2)**. Cytosolic Na^+^ can also inhibit the activity of inward K^+^ channels, but this was only found for the salt-tolerant *Aster tripolium* and not for *A. amellus* (Véry et al. 1998). To our surprise, the maximal conductance of GORK-like channels was hardly affected by the presence of 50 mM Na^+^ in the external medium **(Fig. 3c and d)**, which indicates that quinoa guard cells have an efficient machinery to keep low cytosolic Na^+^ concentrations.

Long-term exposure of quinoa plants to salt in the root medium does affect the conductance of K^+^ channels **(Fig. 3a and b)**, but the impact is moderate and probably caused by indirect effects of salt stress, such as changes in gene transcription. Such indirect regulation mechanisms may also explain the reduced ABA sensitivity of anion channels of plants grown with 500 mM NaCl **(Fig. 4c)**. This reduced conductance is linked to lowered ABA response of guard cell action and stomata movements. However, in control conditions, salt-stressed quinoa plants have stomata with a smaller aperture **(Fig. 5e)**, which matches the lower stomatal conductance of these plants **(Fig. 1a).**

### Moderate ABA responses enhance the water use efficiency of quinoa

Our findings suggest that the moderate ABA responses observed in quinoa may be a key factor contributing to its high water use efficiency under saline conditions. When plants encounter high NaCl concentrations in the soil, quinoa plants initiate stomatal closure, likely mediated by ABA, yet they retain sensitivity to environmental cues such as light and CO_2_. This allows the plants to maintain photosynthesis and growth even under osmotic stress. Rather than triggering a full and irreversible stomatal shutdown, as reported for some glycophytes where ABA leads to prolonged closure (Geilfus et al. 2015), quinoa appears to use ABA signaling in a more nuanced way – acting as a regulator to fine-tune stomatal aperture and water transport dynamics. This finely tuned guard cell behavior enables quinoa to conserve water while sustaining gas exchange—an adaptive trait for thriving in saline soils. By uncovering how halophyte guard cells integrate salt and ABA signals, our study offers a framework for improving crop salt tolerance by targeting traits such as moderated ABA sensitivity and anion channel flexibility—key features for breeding or engineering crops suited to saline and drought-prone environments.

## Acknowledgments

This work was supported by the DFG (Deutsche Forschungs Gemeinschaft) grant SCHE 2148/1-1 to S.S and grant HE 1640/44-1 to RH. M.M. was funded by DFG - TRR 356.

---

**Figure S1.**
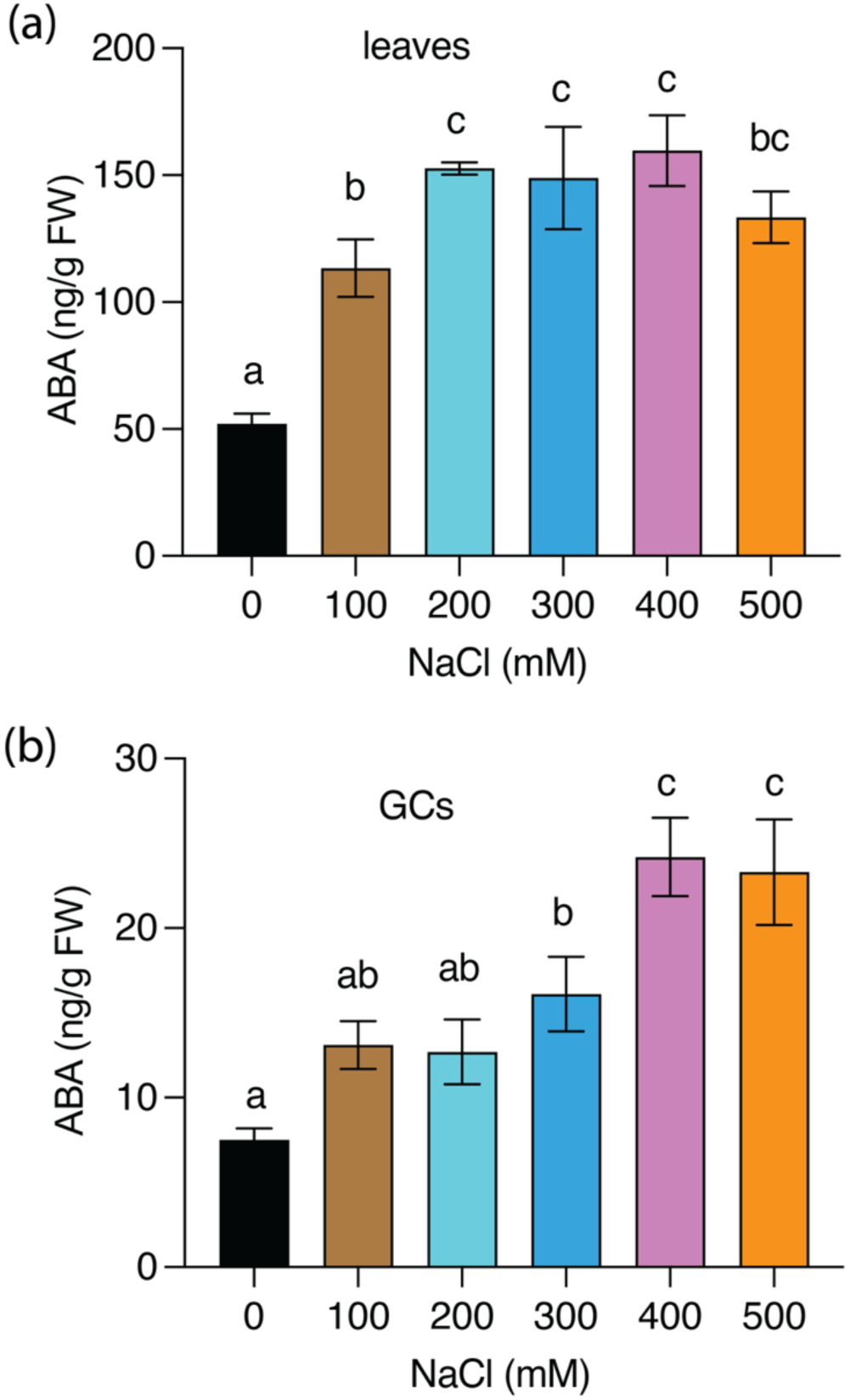
ABA concentrations in quinoa leaves and guard cells. Plants were subjected to long-term salt treatment with varying NaCl concentrations (0, 100, 200, 300, 400, and 500 mM). **(a)** ABA content in leaves. Mean values ± SE of six preparations per treatment. **(b)** ABA content in guard cells. Mean values ± SE of four to five guard cell preparations per treatment. Statistically significant differences were determined by one-way ANOVA followed by Fisher’s LSD post hoc test (see Table S3).

**Figure S2.**
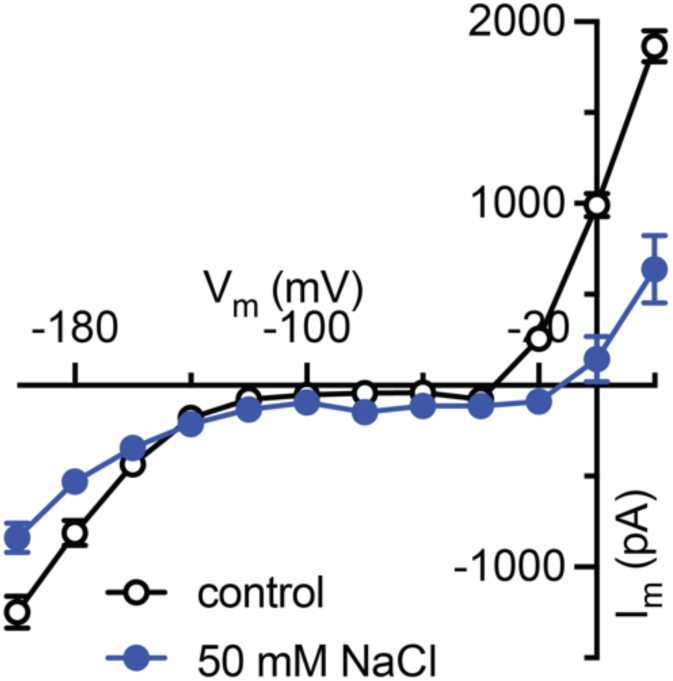
Impact of cytosolic Na^+^ on K^+^ channel activity in quinoa guard cells. Current–voltage relationships of steady-state K^+^ currents recorded in guard cells with or without cytosolic NaCl loading (n = 8).

**Figure S3.**
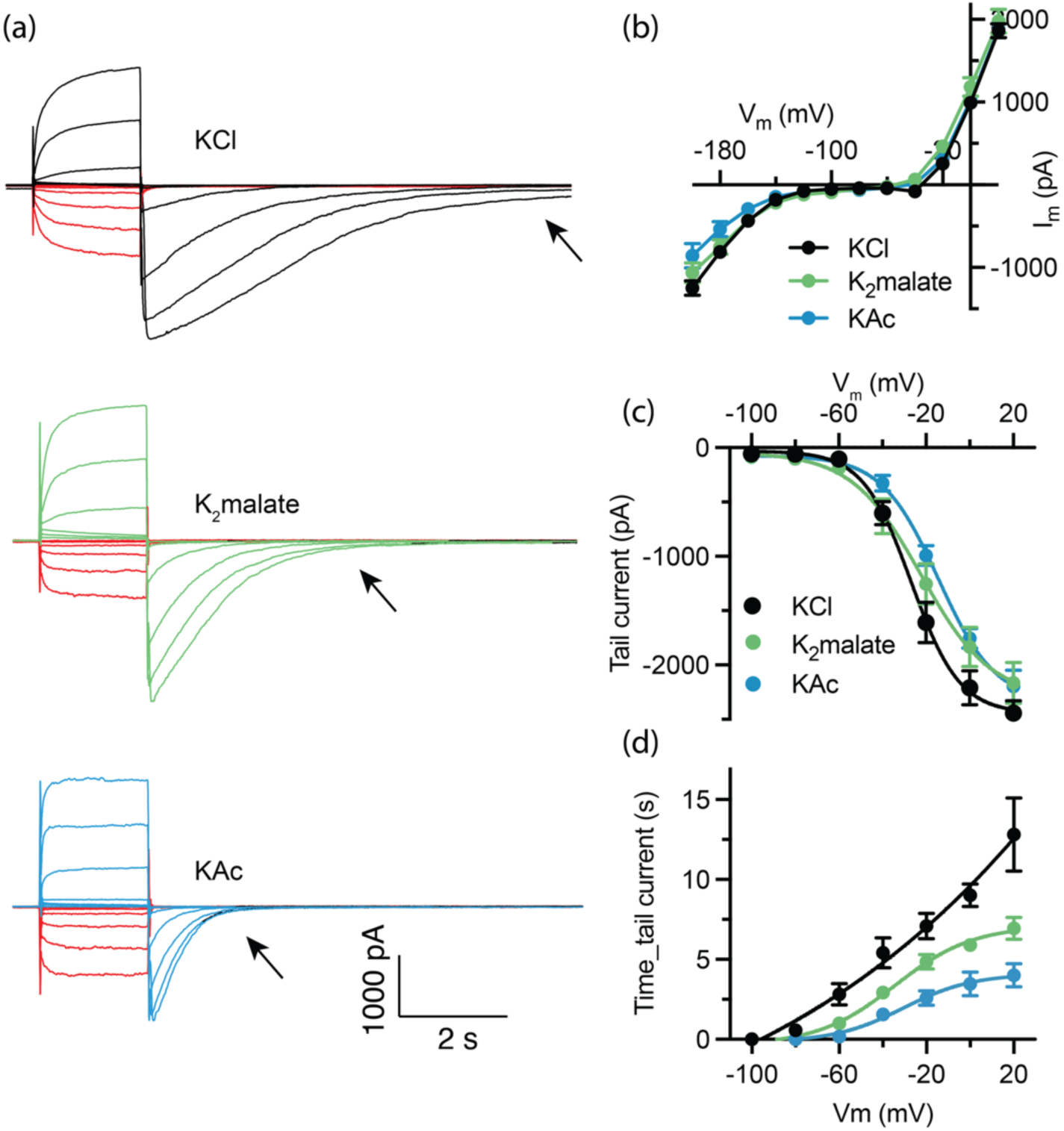
Transient depolarization activates guard cell inward K^+^ and anion channels. **(a)** Representative K^+^ current traces recorded in guard cells with electrodes filled with KCl, K_2_-malate, or K-acetate. Inward currents (red) were activated by hyperpolarizing voltages (–100 to –200 mV), while outward currents (black, green, and blue) were activated by depolarizing voltages (–80 to +20 mV). Arrows indicate tail currents following depolarization. **(b)** Steady-state current–voltage relationships derived from the recordings shown in (a). **(c)** Tail current amplitudes plotted against the membrane potential. **(d)** Duration of tail currents plotted against the membrane potential.

